# LRRC15 mediates an accessory interaction with the SARS-CoV-2 spike protein

**DOI:** 10.1101/2021.09.25.461776

**Authors:** Jarrod Shilts, Thomas W. M. Crozier, Ana Teixeira-Silva, Ildar Gabaev, Edward J. D. Greenwood, Samuel James Watson, Brian M. Ortmann, Christian M. Gawden-Bone, Tekle Pauzaite, Markus Hoffmann, James A. Nathan, Stefan Pöhlmann, Paul J. Lehner, Gavin J. Wright

## Abstract

The interactions between severe acute respiratory syndrome-related coronavirus 2 (SARS-CoV-2) and human host factors enable the virus to propagate infections that lead to COVID-19. The spike protein is the largest structural component of the virus and mediates interactions essential for infection, including with the primary ACE2 receptor. We performed two independent cell-based systematic screens to determine whether there are additional proteins by which the spike protein of SARS-CoV-2 can interact with human cells. We discovered that in addition to ACE2, expression of LRRC15 also causes spike protein binding. This interaction is distinct from other known spike attachment mechanisms such as heparan sulfates or lectin receptors. Measurements of orthologous coronavirus spike proteins implied the interaction was restricted to SARS-CoV-2, suggesting LRRC15 represents a novel class of spike binding interaction. We localized the interaction to the C-terminus of the S1 domain, and showed that LRRC15 shares recognition of the ACE2 receptor binding domain. From analyzing proteomics and single-cell transcriptomics, we identify LRRC15 expression as being common in human lung vasculature cells and fibroblasts. Although infection assays demonstrated that LRRC15 alone is not sufficient to permit viral entry, we present evidence it can modulate infection of human cells. This unexpected interaction merits further investigation to determine how SARS-CoV-2 exploits host LRRC15 and whether it could account for any of the distinctive features of COVID-19.

**In brief:** We present evidence from genome-wide screening that the spike protein of SARS-CoV-2 interacts with human cells expressing LRRC15. The interaction is distinct from previously known classes of spike attachment factors, and appears to have emerged recently within the coronavirus family. Although not sufficient for cell invasion, this interaction can modulate viral infection. Our data point to an unappreciated host factor for SARS-CoV-2, with potential relevance to COVID-19.

**Highlights:**

- Two systematic cell-based screens for SARS-CoV-2 spike protein binding identify LRRC15 as a human host factor
- Interaction with LRRC15 is reproducible in different human cell lines and independent of known glycan or ACE2 binding pathways
- The C-terminal S1 domain of SARS-CoV-2 spike binds LRRC15 with sub-micromolar affinity, while related coronavirus spikes do not
- LRRC15 is expressed in tissues with high ACE2 levels and may modulate infection

## Introduction

Coronaviruses including Severe acute respiratory syndrome coronavirus 2 (SARS-CoV-2) have evolved to recognize host proteins through binding interactions that facilitate viral infection. The ongoing COVID-19 pandemic caused by SARS-CoV-2 has distinguished itself from past coronavirus outbreaks by its virulence and spread (Hu et al., 2020), suggesting that the virus has acquired particularly efficient mechanisms to target its hosts (Gordon et al., 2020; Li, 2015). Interactions involving the viral spike protein are particularly important to characterize, not only because the spike protein enables cell entry by binding to ACE2 (Walls et al., 2020) but also because the spike protein is the central component of many of the current vaccines and therapeutics under development (Harvey et al., 2021; Izda et al., 2021). As the most prominent structural feature of the virion surface, it also is positioned to mediate several key stages of viral pathogenesis, and thus its interactions could hold clues to COVID-19 pathology.

While the spike protein is known to interact with human ACE2 receptor for host cell entry, there has been considerable debate around whether ACE2 alone is sufficient to explain COVID-19 pathology (Puray-Chavez et al., 2021; Sanchez-David et al., 2020; Shahriari Felordi et al., 2021; Zamorano Cuervo and Grandvaux, 2020). For example, ACE2 expression in certain target cells appears to be very low, and other questions remain regarding whether additional factors beyond ACE2 may explain the broad cellular tropism and potent infectivity of SARS-CoV-2 (Hikmet et al., 2020; Osuchowski et al., 2021; Shilts et al., 2021). We therefore employed systematic, unbiased screening approaches to identify additional host proteins which interact with the spike of SARS-CoV-2, and then characterized their activity. Two independent but complementary screening methods identified LRRC15 as a novel host factor inducing binding of the spike protein to human cells by a previously unknown mechanism.

## Results

### Systematic screening for SARS-CoV-2 spike binding interactions identifies LRRC15

While several host proteins have been identified that interact with the spike protein of SARS-CoV-2 based on homology to SARS-CoV-1 (Amraei et al., 2021; Clausen et al., 2020; Hoffmann et al., 2020) or heuristics like the C-end rule (Cantuti-Castelvetri et al., 2020; Daly et al., 2020), there have been few systematic investigations to determine if other cell surface or transmembrane proteins are targeted by the virus. We therefore carried out two independent large-scale screens in separate laboratories to search for additional cellular interaction targets which, when expressed, allow human cells to bind spike protein. In the first screen, we arrayed a library of 2,363 full-length human cDNAs encoding most cell surface membrane proteins in the human genome (Wood and Wright, 2019) and individually transfected them into HEK293 cells (Figure 1A). Transfected cells in each individual well were stained with fluorescently-labelled tetramers of the full SARS-CoV-2 spike extracellular domain, and binding was measured by flow cytometry (Supplemental Figure 1). The second screen used a pooled genome-wide CRISPR activation library in RPE1 cells to identify genes which, when upregulated, induce the binding of the spike protein S1 domain formatted as an Fc fusion construct (Figure 1B).

**Figure 1.**
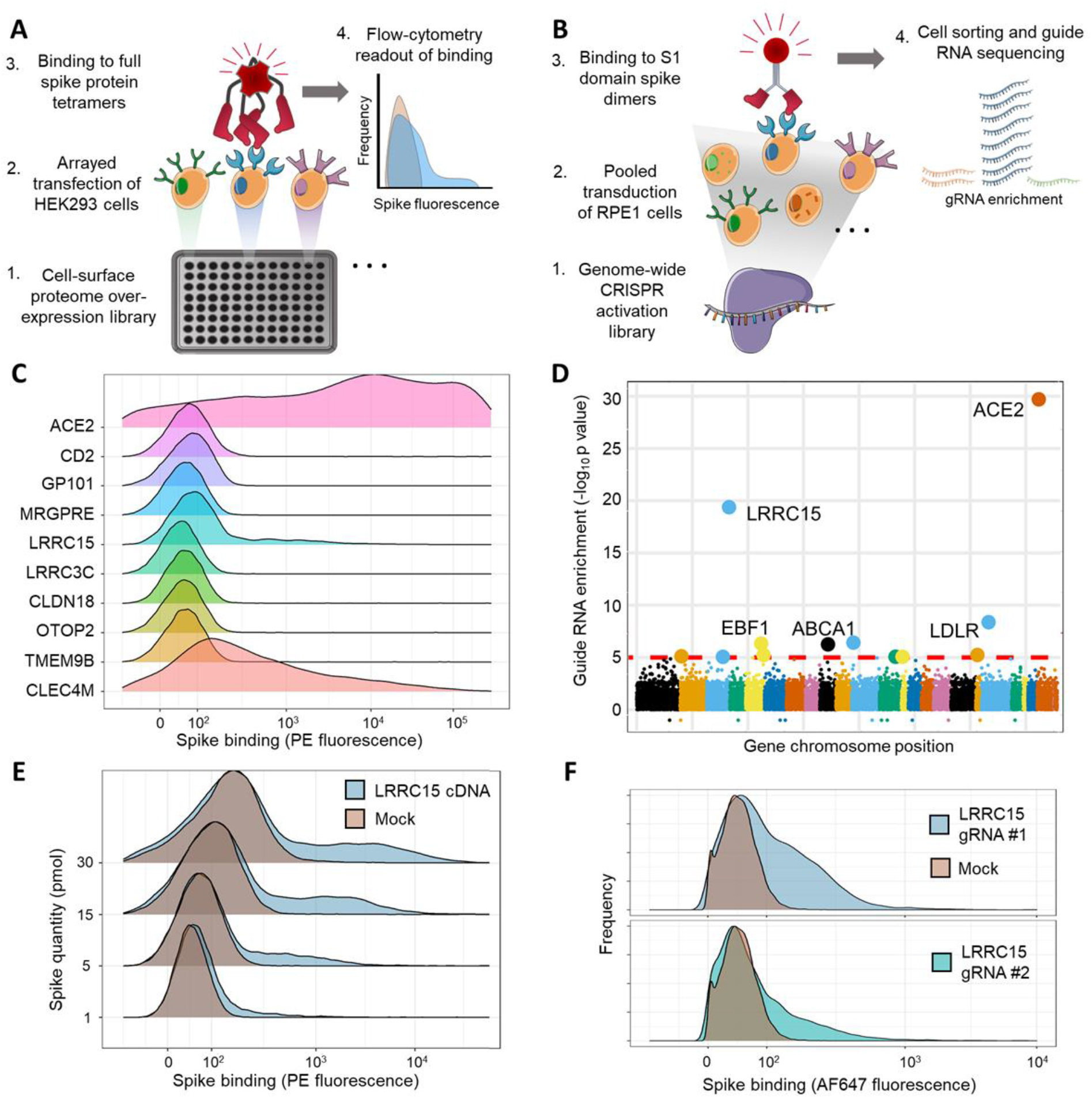
Arrayed transmembrane protein screening and pooled genome-wide CRISPR activation screening identify LRRC15 as binding SARS-CoV-2 spike protein. (A) Schematic of arrayed cell-based screening to identify host factors that cause SARS-CoV-2 spike protein binding. HEK293 cells were transfected in individual wells of microtiter plates with full-length cDNA constructs encompassing a near-comprehensive library of human membrane proteins. Each well in the array was tested for binding to fluorescent tetramers of full-length SARS-CoV-2 spike by flow cytometry. (B) Schematic of pooled CRISPR activation screening. RPE1 cells expressing a SunTag CRISPRa system were transduced with guide RNAs to activate transcription of all genes in the human genome. Cells that bound to Fc protein fusions of the spike S1 domain were sorted by FACS and sequenced to measure guide RNA enrichment. (C) Top-ranked hits from arrayed cDNA screening. Distributions of cell fluorescence by flow cytometry are shown after incubation with fluorescent spike protein tetramers for cDNAs which produced top-ranked signals in the arrayed screen. CD2 is included as a negative control. (D) Guide RNA enrichment of genes promoting SARS-CoV2 spike protein binding using CRISPR activation screening. Genes are ordered according to their positions across the genome and the statistical significance of their respective guide RNA enrichments post-sorting. (E) Independent transfections of LRRC15 cDNA induce spike binding. Flow cytometry traces for HEK293 cells transiently transfected with LRRC15 cDNA compared to mock-transfected controls. Different quantities of full-length spike protein were applied, as indicated along the y-axis. (F) Single guide RNA clones validate *LRRC15* as a gene that induces spike binding. Flow cytometry traces for RPE1 cells transduced and selected for individual LRRC15-activating guide RNAs.

Both screens identified ACE2 as the top-ranked gene as expected, along with other known spike-binding receptors including CLEC4M in the arrayed screen (Figure 1C) and the transcription factor GATA6 in the CRISPR activation screen (Israeli et al., 2021) (Figure 1D). The CRISPRa screen was able to detect even weak spike binding signals, identifying the transcription factor EBF1 despite causing only a small increase in ACE2 levels (Supplemental Figure 2, Supplementary Table 1). Surprisingly, both screens converged on a gene encoding a membrane protein called leucine rich repeat containing protein 15 (LRRC15). The top-ranked genes of both screens were individually verified by either re-arraying cDNAs individually (Figure 1E), or by testing single CRISPR guide RNA clones (Figure 1F). In both cases LRRC15 was confirmed as a host factor that induces spike binding.

### LRRC15 binding to spike protein is specific and reproducible

After discovering the interaction between the spike protein and LRRC15, we sought to validate and characterize the effect of LRRC15. We were intrigued by the spike protein staining profile of cells transfected with LRRC15, where only a subpopulation of cells gained binding according to a long-tailed distribution. Staining with an antibody against the LRRC15 extracellular domain demonstrated that this profile matched LRRC15 cell surface expression (Figure 2A). Subsequently we also wanted to address two common confounding mechanisms that could account for the cell surface binding. First, we excluded whether LRRC15 is non-specifically binding to streptavidin or other molecules in our staining reagents, finding that cells overexpressing LRRC15 did not bind control protein tetramers of Cd4-based linker tags (Figure 2B, left). Second, we tested whether upregulation of LRRC15 triggers binding through heparan sulfate proteoglycans, which are known to broadly bind many proteins (Cagno et al., 2019; Sharma et al., 2018) including SARS-CoV-2 spike (Clausen et al., 2020). We observed that genetic ablation of surface heparan sulfates did not affect spike binding to cells presenting LRRC15, indicating little or no contribution of heparan sulfates (Figure 2B, right). We then tested if alternate LRRC15 overexpression methods would also cause a gain of spike protein binding in a more physiologically relevant cell type separate from the initial screens: CaLu-3 lung epithelial cells. Different lentivirally-delivered LRRC15 constructs or CRISPR activation again consistently caused cells to gain spike binding (Figure 2C).

**Figure 2.**
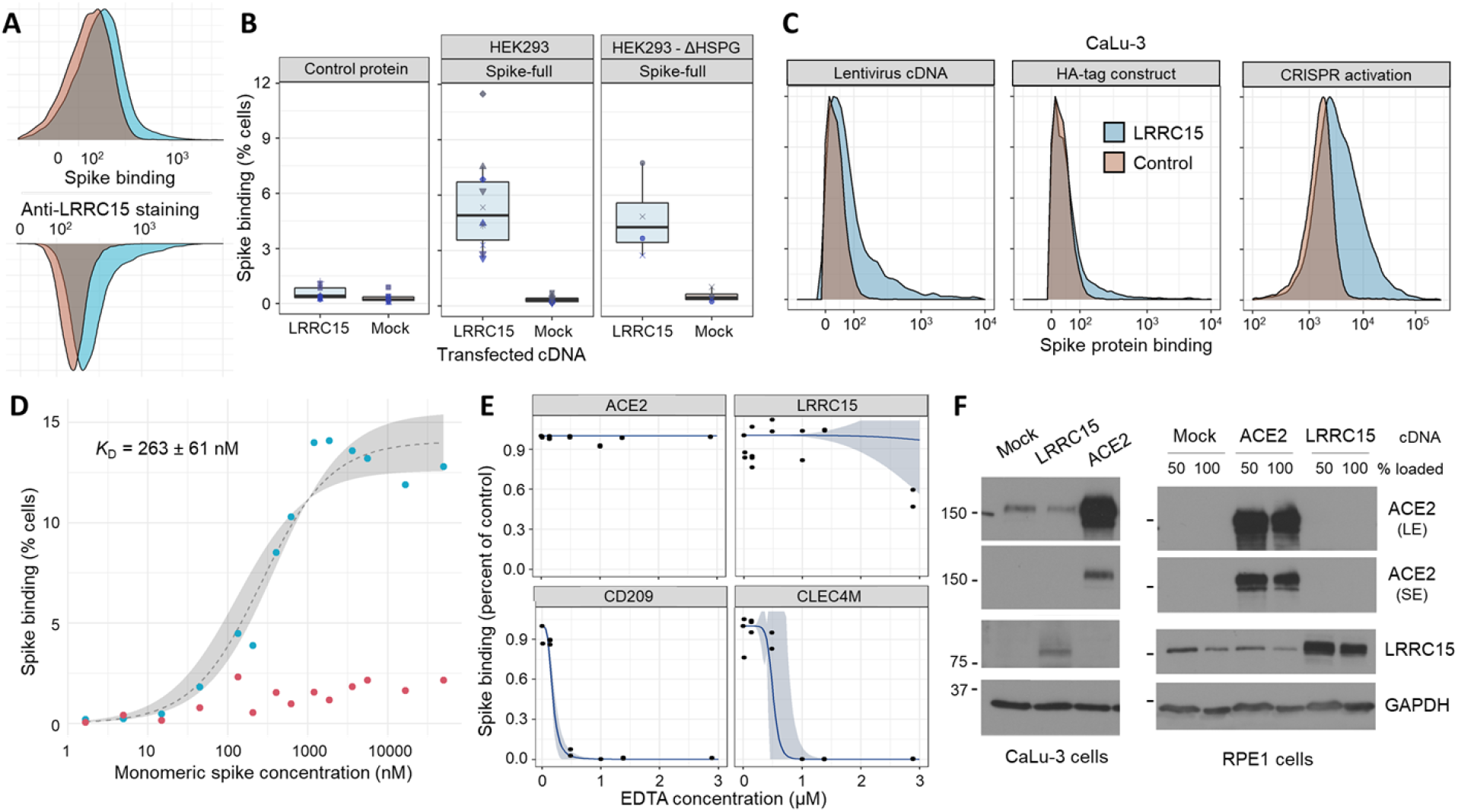
The spike:LRRC15 interaction is robust to cellular context and differs from previously described spike-binding receptors. (A) Comparison of the cell surface staining profiles of HEK293 cells transfected with LRRC15 cDNA withspike protein (top) and anti-LRRC15 antibody (bottom). (B) LRRC15 binding to spike is specific and independent of heparan sulfate proteoglycans. Boxplots summarizing spike binding of LRRC15-transfected cells compared to mock-transfected cells in wild-type HEK293 cells (center panel), a HEK293 strain deficient in cell surface heparan sulfation following SLC35B2 knockout (right panel) and HEK293 cells binding a control tetramer instead of spike (left panel). (C) LRRC15 expression consistently induces spike binding. LRRC15 overexpression was achieved in a CaLu-3 human lung cell line using three different approaches and increases in spike binding as quantified by flow cytometry. (D) Binding of SARS-CoV-2 spike protein to LRRC15-expressing cells is saturable. Spike binding to LRRC15-expressing (blue) and control (red) cells was quantified by flow cytometry over a wide range of fluorescently-conjugated monomeric spike concentrations. A sigmoidal regression curve was fit (grey) to estimate the equilibrium dissociation constant. (E) EDTA blocks known lectin receptors for spike protein but does not prevent LRRC15 binding. Dose-response curves (blue) are fit to flow cytometry measurements of spike binding to HEK293 cells under different concentrations of EDTA. (F) Expression of LRRC15 is not linked to ACE2 protein production. Western blots with antibodies against human ACE2 and LRRC15 detect no upregulation of ACE2 when LRRC15 is overexpressed or vice versa in either an ACE2-negative cell line (RPE1) or ACE2-positive cell line (CaLu-3). ACE2 is shown as both short (SE) and long (LE) exposures.

To further test the specificity of the interaction, we evaluated two other essential criteria for demonstrating an interaction, which are that the binding be saturable and have sufficient affinity to plausibly occur under physiologic conditions. We covalently conjugated the monomeric spike S1 domain to a fluorophore and measured binding to LRRC15-expressing cells over a wide range of spike concentrations (Figure 2D). We observed binding that was saturable and, by fitting a dissociation curve to the data, we could estimate the monomeric affinity as approximately 260 ± 60 nM in terms of its dissociation constant (p = 0.001, t = 4.331). While weaker than the extremely strong spike-ACE2 interaction, which has a dissociation constant in the range of 1-10 nM (Lan et al., 2020; Walls et al., 2020; Wrapp et al., 2020) (Supplemental figure 3), it is within an order of magnitude of the SARS-CoV-1 spike binding interaction with ACE2 and similar to other known viral receptor interactions (Shang et al., 2020; Wang, 2002).

### Prior known viral receptors do not account for spike binding upon LRRC15 expression

Although these features appear to distinguish LRRC15 from previously-described receptors for SARS-CoV-2, we wanted to investigate the possibility that this apparent interaction was due to LRRC15 overexpression causing another spike-binding receptor to become active. The two principal classes of known spike-binding receptors are the protein ACE2 and C-type lectins (including CD209, CLEC4M, CD207, ASGR1 (Anisul et al., 2021; Gao et al., 2020; Hoffmann et al., 2021)). NRP1 binding to spike is dependent on proteolytic processing by furin and thus was not expected in our screens (Daly et al., 2020). Binding to C-type lectin family receptors is cation-dependent (Kolatkar and Weis, 1996; Lee et al., 2011) and so will be ablated by cation chelating agents such as EDTA. We observed that titrating EDTA at concentrations above those required to block interactions with two control C-type lectin receptors had no effect on the LRRC15 interaction with spike protein (Figure 2E). Similarly, human cells overexpressing LRRC15 had no detectable increase in ACE2 protein (Figure 2F) or transcript levels (Supplemental Figure 4), further supporting the distinctness of the LRRC15 interaction.

### LRRC15 - spike interaction is distinct to SARS-CoV-2

SARS-CoV-2 is closely related to other coronaviruses including SARS-CoV-1, which was responsible for the SARS epidemic. Although the majority of their basic biology is shared between viruses (V’kovski et al., 2021), differences do exist which are important in accounting for why COVID-19 has reached levels of global morbidity and mortality not seen with previous coronavirus diseases. We asked if this LRRC15 interaction was conserved across other coronaviruses or distinct to SARS-CoV-2. We selected coronavirus lineages that share ACE2 as their primary receptor, including the closely-related SARS-CoV-1 betacoronavirus and the more distant NL63 alphacoronavirus (Fahmi et al., 2020). Each of the three respective spike protein orthologs were recombinantly expressed and purified as both full-length ectodomains and as the S1 domain only (Figure 3A). All spike proteins bound ACE2-expressing cells as expected. Surprisingly, the gain in spike binding upon LRRC15 expression was seen only with SARS-CoV-2 (Figure 3B). We asked whether this binding pattern may relate to differential glycosylation of the spike proteins which can shield protein interaction surfaces, and indeed observed that enzymatic deglycosylation of the SARS-CoV-1 spike restored LRRC15 binding (Supplemental Figure 5).

**Figure 3.**
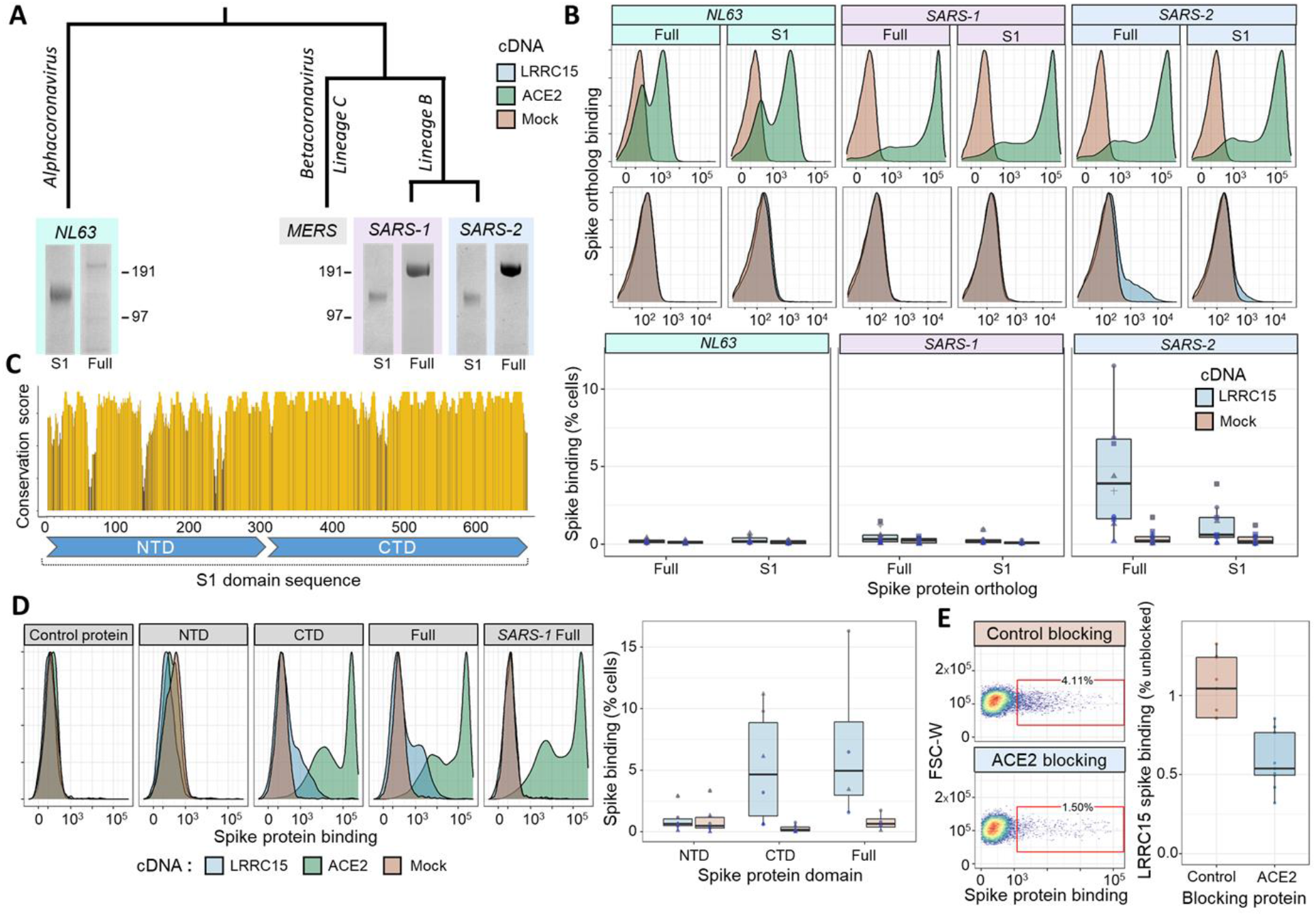
LRRC15 uniquely interacts with the SARS-CoV-2 spike protein and shares a binding interface with ACE2. (A) Recombinant expression of spike proteins from across the coronavirus family. A labeled phylogenetic tree indicates the relative divergences of coronaviruses above Coomassie-stained gel images of purified recombinant spike proteins. For each virus the full extracellular domain was produced along with constructs of only the S1 domain. (B) Orthologous spike proteins bind ACE2 but only SARS-CoV-2 strongly binds LRRC15. Flow cytometry traces of spike proteins binding to HEK293 cells overexpressing ACE2 (green, top) or LRRC15 (blue, bottom) compared to control cells (red). Quantified replicates for LRRC15 binding are displayed as boxplots below. (C) Physical conservation of the amino acids in SARS-CoV-2 spike compared to SARS-CoV-1. The rolling average of physico-chemical conservation scores across the spike S1 domain sequence is indicated. Regions corresponding to the N-terminal domain (NTD) and C-terminal domain (CTD) are annotated below. (D) LRRC15 binding localizes to the spike C-terminal S1 domain. Flow cytometry traces of binding by the CTD but not NTD to LRRC15-transfected cells (left) are shown next to the quantified binding (right). (E) Recombinant ACE2 competitively inhibits spike binding to LRRC15. Dotplots of spike binding by flow cytometry to cells where spike protein was pre-incubated with a control protein or pre-incubated with the ACE2 extracellular domain.

### ACE2 and LRRC15 share a binding domain in the spike C-terminal S1 region

A protein sequence alignment between the spike proteins of SARS-CoV-1 and SARS-CoV-2 shows that although they are broadly conserved, there are some notable differences between regions of the protein (Figure 3C). This led us to ask where the binding to LRRC15 localizes. We separately generated SARS-CoV-2 spike constructs of the N-terminal domain (NTD), which has functions related to glycan recognition and presentation (Li, 2016), and the C-terminal domain (CTD), which includes the ACE2 receptor binding domain. We observed that all binding was accounted for by the spike CTD (Figure 3D). Because ACE2 and LRRC15 shared this binding domain, and the CTD of coronavirus spike proteins are generally known for harboring protein-protein interactions with receptors, we hypothesized that LRRC15 and ACE2 may compete for spike binding. Pre-incubation of spike with recombinant ACE2 did substantially block LRRC15 binding (Figure 3E, p < 0.001, U = 81). These findings suggest a novel function of the receptor-binding region of the SARS-CoV-2 spike protein which appears to be a relatively recent evolutionary innovation.

### LRRC15 is found in mesenchymal and endothelial cells within tissues where ACE2 is present

After accumulating biochemical evidence for this novel interaction, we next sought to understand its potential role in COVID-19 pathology. There is little prior literature on LRRC15, so we began by establishing where in the human body this protein is present, and how that distribution matches with the known tropism of SARS-CoV-2. We examined a detailed whole-body proteomic atlas (Wang et al., 2019) for evidence of LRRC15 expression. We compared the LRRC15 tissue distribution to ACE2, which, as the viruses’ primary receptor, should mark which tissues are virally susceptible (Figure 4A). Despite emerging independently from a genome-wide screen, LRRC15 shared similarities to the tissue distribution of ACE2, with major targets of viral infection such as the lungs and gastrointestinal tract also showing the most abundant expression of LRRC15 (Figure 4B).

**Figure 4.**
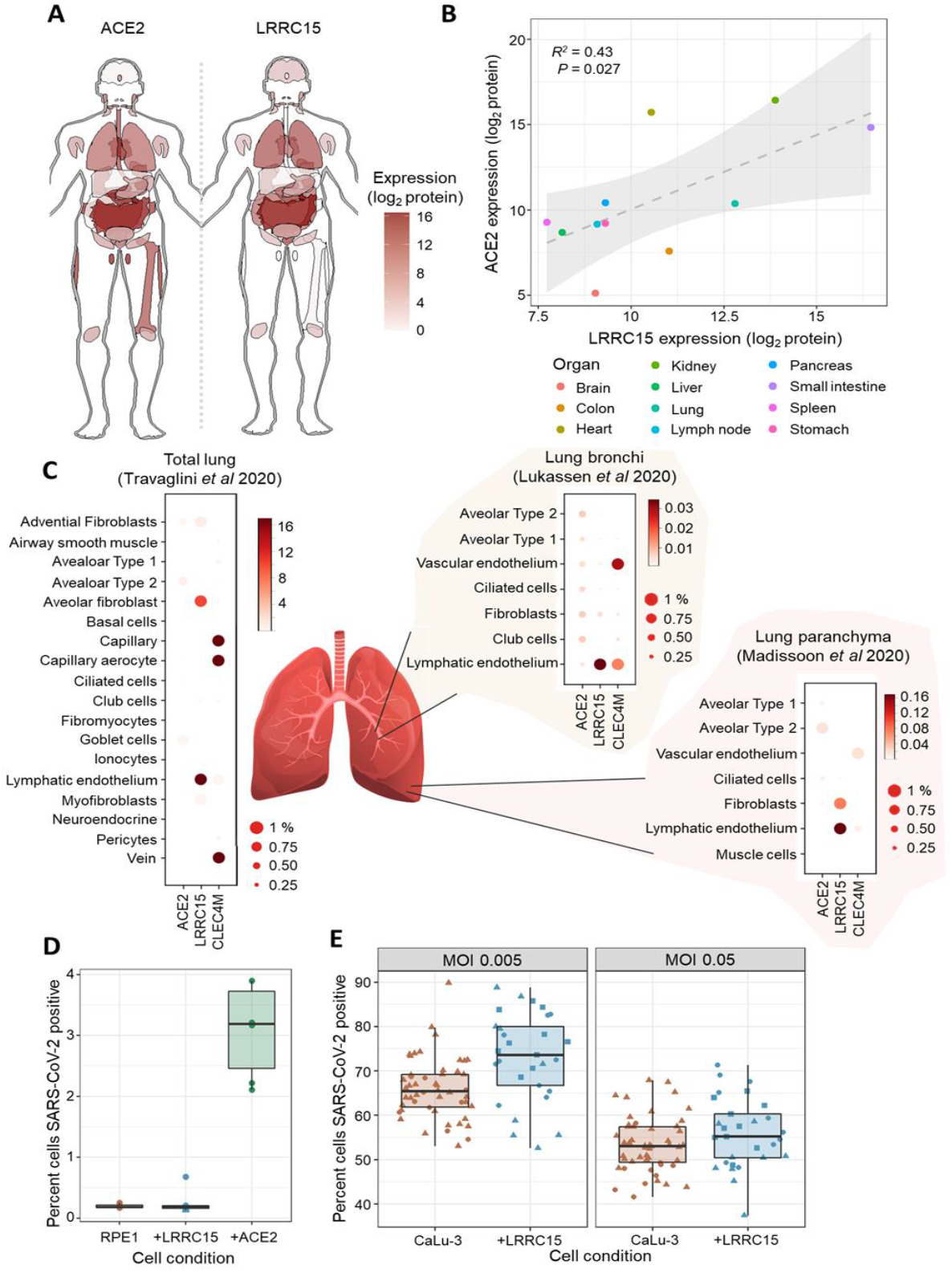
LRRC15 is expressed in virus-susceptible tissues and can modulate SARS-CoV-2 infection. (A) Tissue distribution of LRRC15 and ACE2 expression based on a whole-body proteomic atlas (Wang et al., 2019). For each tissue, protein expression is reflected by the red color intensity scale provided. (B) Organs with high ACE2 expression tend to also express substantial LRRC15. Each data point represents protein abundance in a major human organ or tissue as measured by mass spectrometry. A linear regression line and 95% compatibility interval is shaded in grey. (C) Single-cell transcriptome measurements of human lung specimens identify cell type distributions for LRRC15 expression. The percentage of cells where a gene transcript was detected is indicated by size, while average counts per cell are color-shaded. Expression of spike receptors *ACE2* and *CLEC4M* are shown for comparison. (D) Expression of LRRC15 is insufficient to make cells permissive to SARS-CoV-2 infection. LRRC15 or ACE2 were overexpressed by transducing RPE1 cells with CRISPRa sgRNA followed by infection with a recombinant SARS-CoV-2-ZsGreen reporter virus. (E) LRRC15 overexpression changes the susceptibility of CaLu-3 lung cells to viral infection. CaLu-3 cells were transduced with lentiviruses to overexpress LRRC15 followed by infection with SARS-CoV-2-ZsGreen. Viral infection was quantified by counting GFP-positive cells by fluorescent microscopy in both (D) and (E).

Because lung tissue is the main source of severe COVID-19 symptoms, and expresses among the highest levels of LRRC15 in the body, we conducted a more detailed analysis of LRRC15 expression within the lungs using public single-cell transcriptomic datasets (Lukassen et al., 2020; Madissoon et al., 2019; Travaglini et al., 2020). Although datasets of the upper respiratory tract did not report LRRC15 either because of lower sensitivity or a true absence (Chua et al., 2020; Vieira Braga et al., 2019), in the lower respiratory tract it was readily detected, predominantly in mesenchymal cells and endothelium (Figure 4C). The overlap with cell types expressing ACE2 was relatively weak, although as previously reported these methods only detect ACE2 at very low levels and incompletely (Hikmet et al., 2020; Sungnak et al., 2020; Wang et al., 2021).

### Cell surface LRRC15 is not sufficient to make cells permissive to SARS-CoV-2 infection

Because there has been only relatively limited prior characterization of LRRC15, it raised ambiguities as to whether it can appear on the surface of cells that express it, or if it acts intracellularly and thus more likely to be involved in stages following viral entry (Satoh et al., 2002; Uhlén et al., 2015). We initially tested whether cell surface LRRC15 could act as an entry receptor for the virus, following our observation that LRRC15 predominantly localizes to the cell surface of our cell line models (Supplemental Figure 6). We challenged RPE1 cells with a recombinant SARS-CoV-2-ZsGreen reporter virus (Rihn et al., 2021). RPE1 cells are not naturally permissive to infection, but become infectable upon introduction of ACE2. However, expression of LRRC15 was insufficient to make the cells permissive to infection (Figure 4D).

### Presence of LRRC15 modulates the ability of SARS-CoV-2 to infect host cells

While entry receptors comprise only one class of host-virus interaction, we next sought to gain a broader view of what functional role LRRC15 has in SARS-CoV-2 infection. In order to examine whether LRRC5 might modulate the efficiency of viral entry into ACE2 positive cells, we employed the naturally susceptible human lung cell line (CaLu-3) that is commonly used as a model for coronavirus infection (Chu et al., 2020). We introduced LRRC15 ectopically to these cells to emulate the expression seen in other human lung cell populations we identified in our expression analysis. Compared to unmodified CaLu-3 cells that lack any detectable LRRC15, the expression of LRRC15 modestly enhanced viral infection when the cells were exposed to moderate titers of virus (Figure 4E, MOI 0.005: p = 0.0008, U = 396; MOI 0.05: p = 0.14, U = 580). While these data suggest LRRC15 may have a functional role in SARS-CoV-2 infection, future experiments in primary cells could reveal the precise pathways where the LRRC15-spike interaction acts, and the extent to which it affects infection in different endogenous contexts.

## Discussion

In this report, we describe LRRC15 as a novel interaction target for the SARS-CoV-2 spike protein. We demonstrate that the binding differs from previously known classes of spike binding receptors, and is distinct to SARS-CoV-2. Although cell surface LRRC15 appears to be insufficient to permit viral entry, we present initial evidence that LRRC15 could modulate a host cell’s susceptibility to SARS-CoV-2. Our analysis of public expression datasets suggests LRRC15 is expressed in tissues susceptible to infection at levels exceeding those of ACE2. In the lung, these include several cell types which have been detected as virally-infected targets in human patient samples (Colmenero et al., 2020; Delorey et al., 2021; Deshmukh et al., 2021; Ramos da Silva et al., 2021; Rendeiro et al., 2021), although interestingly the highest LRRC15 levels are not associated with the alveolar epithelial cell types that are most commonly studied. While the fibroblast and endothelial populations where LRRC15 is most abundant have clear significance to COVID-19 pathology (John et al., 2021; Kondo et al., 2021; Melms et al., 2021; Rendeiro et al., 2021), it will be important to determine how the lower level of direct viral infection seen in these cell populations may relate to LRRC15’s endogenous role in infection, or whether cells outside of the lung can reveal where LRRC15 expression is most clinically relevant. For example, case reports of SARS-CoV-2 infection in the brain implicate lymphatic vessel cells where LRRC15 is particularly plentiful (Bostanciklioğlu, 2020). Notably, LRRC15 was found to be significantly upregulated in the lungs of fatal COVID-19 patients compared to controls (Wu et al., 2020), which further emphasizes the value of closer investigation into LRRC15 in COVID-19 now that our work adds a potential biochemical mechanism.

The normal molecular function of LRRC15 in the human body is not well characterized (Dolan et al., 2007; Purcell et al., 2018; Satoh et al., 2002). The protein’s 15 leucine-rich repeat domains in its extracellular region suggest that it may engage in protein-protein interactions (analogous to LRRC15’s closest human paralog, the platelet receptor member GP5), although it contains no obvious cytoplasmic signaling motifs (Ben-Ami et al., 2020). Similarly, it may also have physical interactions with components of the extracellular matrix (Satoh et al., 2005). Although LRRC15 has been found by us and previous researchers to be abundant on the cell surface, some quantity appears present in intracellular membranes, and these could conceivably be acting through other pathways beneficial to the virus’ proliferation such as via directing protein trafficking (O’Prey et al., 2008). Based on our inability to recombinantly produce a functional version of LRRC15 when truncating its extracellular domain, we conjecture that the transmembrane domain or cytoplasmic cysteine-rich region of the protein may also be required. A recent study on LRRC15 similarly found the presence of a membrane-spanning domain was essential for binding activity to be observed (Cao et al., 2021). Despite its lack of biochemical characterization, LRRC15 is remarkably well-conserved among mammals, including being approximately 90% identical in sequence between human and nearly all proposed host species for SARS-CoV-2 (Supplemental figure 7). Elucidating what function LRRC15 has evolved for within these hosts could provide a critical clue to determining how this interaction acts during viral infection.

Because of the wide clinical significance of the SARS-CoV-2 spike protein, awareness of this novel interaction has other implications. Several classes of successful therapeutics operate by blocking spike protein interactions with human host receptors, most notably in the form of ACE2-blocking biologics (Glasgow et al., 2020; Pinto et al., 2020; Zoufaly et al., 2020). Our identification of a shared binding domain between LRRC15 and ACE2 implies many of these may also prevent spike - LRRC15 interactions, at least extracellularly. The development of drugs targeting LRRC15 has already been initiated by oncologists who observed elevated LRRC15 in certain tumors (Dominguez et al., 2020; Purcell et al., 2018). These could provide a means for further dissecting the biology of LRRC15 in infection and eventually may be possible to repurpose in patient settings.

Our investigations so far have only just begun to reveal what roles LRRC15 may have in COVID-19. Characterizing the functional and molecular underpinnings of this interaction may provide useful targets for therapeutics and guide our understanding of what makes SARS-CoV-2 so distinctly pathogenic.

## STAR Methods

### HEK293 cell culturing

For both binding assays and the production of recombinant proteins, suspension HEK293-E cells (Durocher et al., 2002) were grown in Freestyle media (Gibco #12338018) supplemented with 1% (m/v) fetal bovine serum (HyClone #SH30071) at 37°C in 5% CO_2_ and 70% humidity. Cells were transiently transfected 24 hours after being split to a density of 2.5×10^5^ cells per mL (Shilts et al., 2021). For protein production, transfections were performed with 0.5 μg plasmid DNA per mL cells, while for cDNA expression, 1 μg per mL was used. When expressing recombinant proteins that were to be covalently biotinylated, cells were also co-transfected with 40 ng per mL of a BirA expression plasmid (Fairhead and Howarth, 2015) and the cell media was supplemented with an additional 100 μM D-biotin (Sigma #2031). For binding assays, cells were incubated a further 40-48 hours after transfection before being used in experiments. For protein production, the incubation time was 96-120 hours.

### Recombinant protein purification

After HEK293 cells were transfected and allowed to incubate, they were centrifuged at 2000 × g for 20 minutes. Recombinant protein constructs all contained signal peptide secretion sequences, and hence the supernatant fraction was collected for purification. For affinity purification to the protein’s His-tags, nickel-nitrilotriacetic acid (Ni-NTA) resin (Thermo Scientific #88221) was prepared with two washes for 10 minutes each in 25 mM imidazole (Sigma #I2399) phosphate buffer. Supernatant filtered through 0.22 μm filters was then mixed with the Ni-NTA resin and incubated with agitation overnight at 4°C. Samples were washed three times with 25 mM imidazole phosphate buffer (Sigma #I2399), incubating 5 minutes in between washes. Finally proteins were eluted using 200 mM imidazole buffer and resin was separated on polypropylene columns (Qiagen #34924). SARS-CoV-2 S1-Fc protein was purified from serum-free supernatant of transfected 293T cells by Protein A affinity chromatography using hiTrap MabSelect PrismA columns (GE Healthcare) and the Äkta-Pure liquid chromatography system. Experimental replicates all feature different batches of purified protein.

### Protein construct designs

SARS-CoV-2 spike protein constructs were taken from previously-published designs (Hoffmann et al., 2020; Shilts et al., 2021). The full extracellular domain (Q14-K1211) was mutated to remove its polybasic cleavage site (682-685 RRAR to SGAG) and introduce a proline stabilizing mutation (986-987 KV to PP). S1 domain truncations were made at Y674. Orthologous spikes were aligned to SARS-CoV-2 to define similar boundaries. These for SARS-CoV-1 were at S14-L666 (S1 domain) and S14-K1193 (full extracellular), and for NL63 at C19-V745 (S1 domain) and C19-K1294 (full extracellular). The boundary for the SARS-CoV-2 N- and C-terminal domains was between F318 and R319 within the S1 domain (Lan et al., 2020). Recombinant ACE2 for blocking experiments had its extracellular domain defined as M1-S740. All recombinant proteins contained a tag and linker region described previously (Brown and Barclay, 1994; Sun et al., 2012) that includes a 6-His tag for purification, fragment of rat Cd4 for stabilization, and a biotin acceptor peptide for enzymatic monobiotinylation. Protein control constructs consisted solely of this tag region. cDNA constructs for LRRC15 corresponded to the complete sequence (NM_001135057.2) derived from a commercial cDNA library (Origene #SC325217). cDNA for ACE2 (NM_021804.2) was derived from a similar expression plasmid (Geneocopia #EX-U1285-M02). All nucleotide sequences were verified by Sanger sequencing.

### Tetramer normalization ELISAs

To calculate optimal stoichiometries for forming recombinant protein tetramers, competitive ELISAs were performed where biotinylated proteins were titrated against streptavidin. 96-well streptavidin-coated plates (Greiner #655990) were rinsed in 175 uL HEPES-buffered saline (HBS) with 0.1% Tween-20 (HBS-T, Sigma #P2287), then blocked for 1 hour in 2% (m/v) bovine serum albumin (BSA, Sigma #A9647) in HBS. In a separate polystyrene 96-well plate (Greiner #650161), a 2x dilution series of the biotinylated recombinant protein was made in a solution of 2% BSA HBS. To each well of the dilution series,1.5 pmol of fluorescently-conjugated streptavidin (Biolegend #405245 or Biolegend #405237) was added and allowed to incubate for at least 1 hour. Samples were then transferred to the blocked streptavidin-coated plate. Free biotinylated proteins were then allowed to be captured by the plate over at least 45 minutes, before three washes with 150 μL HBS-T. For primary antibody staining, 1.6 μg/mL of a monoclonal OX68 antibody against the recombinant protein tag region was incubated for 1 hour. Following three more HBS-T washes, 0.2 μg/mL of a secondary anti-mouse IgG antibody linked to alkaline phosphatase (Sigma #A9316) was incubated for 30 minutes. Plates were washed a final three times with HBS-T before 60 μL substrate was added in the form of 2 mg/mL para-Nitrophenylphosphate (Sigma #P4744) in diethanolamine buffer. The reaction was allowed to proceed for approximately 30 minutes before absorbance at 405 nm was measured on a Tecan Spark plate reader. The minimum concentration of biotinylated protein at which signal was reduced to baseline (indicating stoichiometric equivalence with the 1.5 pmol streptavidin) was selected for assembling tetramer reagents.

### Cytometry binding assays

Human cell lines transiently transfected or lentivirus-transduced to overexpress LRRC15 (or mock-treated) were transferred into 96-well u-bottom plates (Greiner #650161) at approximately 5×10^4^ cells per well. To 50 μL cells, 50 μL of DAPI was added for a final concentration of 1 μM DAPI. Plates were then incubated on ice for 5 minutes. To remove the supernatant, plates were centrifuged at 4°C for 3 minutes at 200 × g. Cells were resuspended in 100 μL solutions of pre-conjugated recombinant protein in 1% BSA in phosphate buffered-saline (PBS) supplemented with calcium and magnesium ions (Gibco #14040133). To assemble recombinant protein tetramers, a fixed quantity of streptavidin linked to R-phycoerythrin (Biolegend #405245) or AlexaFluor 647 (Biolegend #405237), typically 15 pmol or 30 pmol, was allowed to incubate for at least 1 hour with the covalently biotinylated recombinant proteins at the concentration empirically determined by the tetramer normalization ELISA (see above). For recombinant Fc fusion proteins, purified spike protein was directly added to a final concentration of 16 μg/mL. Cells were left to bind the recombinant proteins over at least 45 minutes on ice. To wash, an additional 100 μL of PBS was added, plates centrifuged again to decant supernatant, then resuspended in 200 μL PBS before another centrifugation. In the case of fluorescent-conjugated tetramer staining, cells were finally resuspended in 1% BSA in PBS, while in the case of Fc fusion protein staining, cells were stained with secondary antibody conjugated to AlexaFluor 647 against human IgG-gamma1 (Jackson #109-605-006), washed, and then resuspended. Cells were measured on a BD LSR Fortessa flow cytometer. The gating strategy is illustrated in the supplementary information.

### Arrayed cDNA screen

The human cDNA library used for the arrayed screen was previously described (Wood and Wright, 2019), and covers the vast majority of human cell surface proteins (Bausch-Fluck et al., 2018, 2015; da Cunha et al., 2009). The screening procedure followed the ‘Cytometry Binding Assays’ protocol above, with the following modifications. After 24 hours of HEK293 cells incubating in a standard shaking flask (Corning #431143), the cells were transferred to u-bottom 96-well plates for transfections. 100 μL of cells at a density of 5×10^5^ cells/mL were then transfected with 200 ng cDNA plasmid. 46 hours after transfection, cells were stained with full-length SARS-CoV-2 spike tetramers around streptavidin R-phycoerythrin (Biolegend #405245) as described above, except only a 5 pmol quantity of protein was used. To avoid screen hits unrelated to protein-protein interactions due to heparan sulfate binding, we used a previously-described HEK293 cell line with a biallelic targeted disruption of the *SLC35B2* sulfate transporter which prevents cell surface heparan sulfate presentation (Gao et al., 2019; Riblett et al., 2016; Sharma et al., 2018). The gate for positive protein staining was set at a fluorescence value of 10^3^ across all plates. All hits from the arrayed screen were repeated individually using the standard procedure described above, which resolved false-positives caused by occasional blockages to the laminar flow of the cytometry instrument when running such a large quantity of samples.

### CRISPRa Screen

A genome-wide CRISPRa screen was performed using a clonal RPE-1 cell line containing SunTag CRISPRa system (a gift from the Tanenbaum lab), where a nuclease-dead Cas9 (dCas9) is fused to 10xGCN4 peptide array (with a P2A-mCherry reporter) together with a GCN4 nanobody fused to GFP and the transactivator VP64. A total of 1.26×10^8^ cells were transduced with CRISPRa sgRNA library (Addgene #1000000091) lentivirus at an MOI of 0.3 (180-fold coverage), which was assessed by flow cytometry (BFP^+^) at 72 hours post infection. sgRNA containing cells were enriched by selecting with puromycin (10 μg/mL) up until the first sort. Cells were harvested for fluorescence activated cell sorting (FACS) at day 10 post-transduction. Trypsinized cells were blocked with rabbit IgG (EMD Millipore, 20 μg/mL) for 10 minutes in sort buffer (PBS + 2% FCS), to which purified S1-Fc (final concentration 16 μg/mL) was added and incubated for 40 minutes on ice. Cells were then washed in sort buffer and stained with an AF647 conjugated F(ab’)2 antibody against Human IgG-gamma1 (Jackson #109-605-006, 1.5 μg/mL) for 30 minutes on ice. A total of 1×10^8^ cells (90% BFP^+^) were sorted on BD-Influx cell sorters. The top 2.8% most AF647^+^ population, amounting to 2.6×10^6^ cells, were collected and used directly for DNA extraction (Qiagen, Gentra Puregene). Cells were gated to ensure mCherry and GFP reporters did not change and sgRNA expression was selected by sorting for BFP^+^ cells. An unsorted library population was maintained separately at 180-fold coverage throughout the experiment, and DNA from this population was also extracted at day 10. sgRNA sequences were amplified and Illumina sequencing adaptors added by two sequential rounds of PCR followed by PCR purification (AMPure XP, Beckman Coulter). Next-generation sequencing was performed on a MiniSeq System (Illumina) using a custom primer. For data analysis, single-end 35 bp reads were trimmed down to the variable sgRNA segment using FASTX-Toolkit and aligned to an index of all sequences in the library using Bowtie 2. Read count statistics were generated using the RSA algorithm (König et al., 2007). Sequencing data have been submitted to SRA under the accession PRJNA762706.

### Antibody staining

For antibody staining for flow cytometry, HEK293 cells were incubated with polyclonal against the LRRC15 extracellular domain (LSBio #LS-C165855) in a 1:50 dilution in 1% BSA PBS. Staining was done in u-bottom 96-well plates (Greiner #650161) for 30 minutes on ice. Plates were then centrifuged at 200 × g for 3 minutes to remove the supernatant and washed twice in PBS. A 1:800 dilution of anti-rabbit IgG secondary antibody conjugated to AlexaFluor 488 (Jackson #111-547-003) in 1% BSA PBS was incubated for 20 minutes on ice. One additional PBS wash was performed before resuspending the cells in 1% BSA PBS and measuring them on a BD LSR Fortessa flow cytometer. For immunofluorescence staining, CaLu-3 were plated on 13 mm, round, #1.5 coverslips (VWR) and allowed to adhere for 72 h. Media was exchanged for methanol free 4% PFA in PBS and incubated for 15 minutes. Coverslips were washed three times in Tris-buffered saline pH 7.4 (TBS) and subsequently permeabilized in 0.1% Triton X-100 in TBS for five minutes before being blocked with 2% BSA (Arcos) in TBS for 20 minutes. Coverslips were then incubated in TBS + 2% BSA containing the indicated dilution of primary rabbit anti-human LRRC15 antibody (ab150376) for 1 h at room temperature, washed three times in TBS + 2% BSA, incubated for 30 minutes in TBS + 2% BSA containing highly cross-adsorbed goat anti-rabbit Alexa Fluor 555 conjugated secondary antibody (2 μg/mL) and DAPI (0.2 μg/mL), then washed three times in TBS and once in distilled water prior to mounting slides with Prolong Glass (Thermo Fisher) followed by overnight curing. Samples were observed on a Zeiss 980 equipped with an Airyscan2 using the 405 nm (DAPI) and 555 (Alexa Fluor 555) lasers in Airyscan mode to provide high resolution images.

### Affinity measurement assay

Cell-binding assays to calculate monomeric affinity were modified versions of ‘Cytometry Cell Binding’ protocol above. Recombinant SARS-CoV-2 spike protein S1 C-terminal domain was dialyzed into PBS (Millipore #71505) and then covalently conjugated to R-phycoerythrin using a commercial amine coupling kit (Abcam #ab102918). Transfected HEK293 cells were transferred to v-bottom 96-well plates (Greiner #651261). Comparatively small quantities of cells were used (10 μL, or about a few thousand cells per well) in order to ensure that the assumptions around free ligand concentration for the 1:1 binding model equation were not violated (Hunter and Cochran, 2016). After the initial wash and DAPI stain, cells were resuspended in a 3x dilution series of spike protein going across the wells of the plate, starting at 16.7 μM diluted in 1% BSA in PBS with calcium and magnesium ions (Gibco #14040133). Binding was allowed to reach equilibrium over an hour at 4°C before two washes in PBS. For washes, cells were centrifuged at 300 × g for 7 minutes at 4°C then supernatant was carefully aspirated by pipette. Cells were finally resuspended in 40 μL 1% BSA PBS and measured with a BD LSR flow cytometer.

### Coomassie protein staining

Purified protein samples were characterized by Coomassie total-protein staining. Proteins were first denatured in NuPAGE sample buffer (Invitrogen #NP0007 and #NP0004) by heating to 70°C for 10 minutes. These were then loaded on to 4–12% gradient Bis–Tris gels (Invitrogen #NP0329) and electrophoresed for 50 minutes at 200 volts. Gels were removed from their cassettes, briefly rinsed in pure water, then stained with Coomassie R-250 (Abcam #ISB1L) overnight at room temperature. After a brief rinse in water, gels were imaged under visible light filters with an Azure c600 system.

### Western blotting

Cells were collected by trypsinization and washed 3 times with PBS (1000 × g, 5 minutes, 4°C). Cell pellets were resuspended in lysis buffer (1% (w/v) digitonin, 1 × Roche cOmplete protease inhibitor, 0.5 mM PMSF, 10 mM Tris-HCL pH 7.4) and incubated on ice for 40 minutes. The lysates were then centrifuged (17,000 × g, 10 minutes, 4°C) and the post-nuclear fractions were transferred to new tubes. The protein concentration of each sample was determined by Bradford assay. Samples were adjusted with TBS buffer and 6 × Laemmli buffer + 100 mM dithiothreitol (DTT) and heated at 50°C for 10 minutes. Samples were separated by SDS-PAGE and transferred to PVDF membranes (Merck), then blocked in 5% milk + PBS-T (PBS + 0.2% (v/v) Tween-20) for 1 hour. Blocked membranes were incubated with primary antibody (LRRC15 [LSBio aa393-422], ACE2 [Abcam Ab108252], GAPDH [GeneTex GTX627408]) in 5% milk +PBST (PBS + 0.2% (v/v) Tween-20) at 4°C overnight, then incubated with peroxidase (HRP)-conjugated secondary antibodies (Peroxidase AffiniPure Goat Anti-Rabbit IgG (H+L) [Jackson ImmunoResearch 111-035-144], Peroxidase AffiniPure Goat Anti-Mouse IgG (H+L) [Jackson ImmunoResearch 115-035-146]) for 90 minutes at room temperature.

### qPCR

Total RNA was extracted using the RNeasy Plus minikit (Qiagen) following the manufacturer’s instructions and then reverse transcribed using Protoscript II Reverse Transcriptase (NEB). Template cDNA (20 ng) was amplified using the ABI 7900HT Real-Time PCR system (Applied Biotechnology or Quantstudio 7, Thermo Scientific). Transcript levels of genes were normalized to a reference index of a housekeeping gene (β-actin).

### Phylogenetics analysis

Homologs for LRRC15 were identified by BLAST searches against the human LRRC15 amino acid sequence, filtering for species reported to be potential SARS-CoV-2 hosts (Liu et al., 2021; Mahdy et al., 2020). Multiple sequence alignments were done by the Clustal Omega algorithm using default parameters (Madeira et al., 2019). Phylogenetic trees were drawn using Jalview (v. 2.11.1.4) from average distance calculations based on the BLOSUM62 substitution matrix. For coronavirus spike protein comparisons, SARS-CoV-1 spike and SARS-CoV-2 spike protein sequences were pairwise-aligned and conservation at each amino acid position was calculated based on an established physico-chemical conservation score (Livingstone and Barton, 1993). Measurements for the alpha and beta coronavirus phylogenetic tree were taken from a prior study on whole-genome nucleotide sequences (Fahmi et al., 2020).

### Expression data analysis

Whole-body proteomics data was downloaded from a prior study (Wang et al., 2019). The data were visualized with the gganatogram pacakge (v. 1.1.1) in R (v. 4.0.3). An expression value of at least 500 molecules (based on intensity-based absolute quantification (Schwanhäusser et al., 2011)) was required for a protein to be displayed as expressed in a given tissue. Lung single-cell RNA sequencing datasets were downloaded from the COVID-19 Cell Atlas and related resources (Lukassen et al., 2020; Madissoon et al., 2019; Sungnak et al., 2020; Travaglini et al., 2020). The data were visualized with the Scanpy package (v. 1.4.5) in Python (v. 3.7.4). Datasets with no detected read counts for either ACE2 or LRRC15 were excluded.

### Statistical calculations

Whenever two groups were compared, a Mann-Whitney U test was used to compare distributions without invoking normality assumptions. Redundant siRNA activity (RSA) analysis was used to calculate p-values to identify statistically significantly enriched genes, comparing read counts from individual sgRNAs from harvested sorted cells to an unsorted library population (König et al., 2007). The statistical significance of the dissociation constant fit in the monomeric binding affinity model was computed using the *nls* function in the R stats package (v. 4.0.3). Correlations are reported as Pearson linear regression coefficients. Boxplots follow the standard Tukey style and illustrate the 25^th^ to 75^th^ percentiles, with the median denoted by the center horizontal line and whiskers showing points within 1.5 times the interquartile range. Different experimental batches are shown as different shapes on plots where all data points are illustrated.

### Generation of cell lines and culture conditions

Human embryonic kidney (HEK-293T), lung adenocarcinoma (CaLu-3), and retinal pigment epithelium (hTERT RPE-1) cells were used in this study. HEK-293T and hTERT RPE-1 cells were grown in Dulbecco Modified Eagle Media (DMEM) supplemented with 10% fetal calf serum (FCS). Calu-3 cells were grown in Minimum Essential Medium (MEM) supplemented with 10% FCS, 2mM GlutaMAX, 1mM Sodium Pyruvate, and non-essential amino-acids. All cell lines were maintained at 37°C and 5% CO_2_. To generate CaLu-3 CRISPRa cells, CaLu-3 cells were transduced with pHRSIN-P_SFFV_-dCas9-VPR-P_SV40_-Blast^R^ and stable integrants were selected with Blasticidin.

### Vector cloning and lentiviral production

CRISPRa constructs and whole genome guide library were created by the Weissmann lab and provided to us by the Tanenbaum lab. Individual CRISPRa guides were cloned into pCRISPRia-v2 (Addgene #84832). ACE2 and LRRC15 cDNAs were cloned into pHRSIN-cSGW vectors expressing puromycin, blasticidin or hygromycin resistance cassettes driven by a pGK promoter. VSV-G lentiviruses were produced by transfection of HEK293T cells with a lentiviral expression vector and packaging vectors pCMVΔR8.91 and pMD.G at a DNA ratio of 3:2:1 using TranslT-293 (Mirus) following the manufacturers recommendation.

### Production of SARS-CoV-2 viral stocks

A total of 1 μg of pCCI-4K-SARS-CoV-2-ZsGreen plasmid DNA and 3 μL of Lipofectamine LTX with 3 μL of PLUS reagent in 100 μL optiMEM were used to transfect BHK-21 cells (ECACC) in 6-well plates. Three days post-transfection, supernatant was transferred to CaLu-3 cultures in 6-well plates. Virus was allowed to propagate in CaLu-3 cells until clear cytopathic effect was observed, around 72 h post infection. Virus containing supernatant was then utilised to seed larger cultures of CaLu-3 cells in T25 flasks to produce viral stocks utilised for infection assays. Viral titre was determined by 50% tissue culture infectious dose (TCID50) in CaLu-3 cells.

### SARS-CoV-2 microscopy infection assays

RPE-1 or CaLu-3 cell lines were seeded into CellCarrier-96 Black optically clear bottom plates (Perkin Elmer) at a density of either ×1×10^4^ cells/well 24 h prior to infection or ×5×10^4^ cells/well 72 h prior to infection respectively. Cells were infected with SARS-CoV-2-ZsGreen (Rihn et al., 2021)at indicated MOIs and infection allowed to proceed for 48 h followed by fixation by submerging plates in 4% formaldehyde for 15 minutes. Cells were stained with DAPI (Cell Signalling Technology, 0.1 μg/mL) at room temperature for 15 minutes prior to washing in PBS and imaging. Images were acquired using an ArrayScan XTI high-throughput screening microscope (ThermoFisher) with a 10x magnification, using the 386 nm or 485 nm excitation/emission filter to detect DAPI and ZsGreen signal respectively across 16-fields per well. HCS Studio software (ThermoFisher) was utilized to analyze images using the Target Activation application. Individual cells were identified by applying masks based on DAPI intensity, excluding small and large objects based on average nuclei size. The DAPI generated nuclei masks were applied to the ZsGreen channel, providing a fluorescent signal intensity of SARS-CoV-2-ZsGreen detected in each cell. These data were exported as .fcs files and analysed in FlowJo. Uninfected control wells were utilised to determine the fluorescence threshold to define SARS-CoV-2 infected ZsGreen^+^ cells and the percentage of infected cells per well across all acquired data.

## Acknowledgements

An RPE-1 cell line expressing dCas9-SunTag_10×_v4_-P2A-mCherry and scFv-GCN4-GFP-VP64 was provided by Marvin Tanenbaum and Jonathan Weissman. pCCI-4K-SARS-CoV-2-ZsGreen was a gift from Professor Sam Wilson, University of Glasgow. FACS experiments were enabled by Dr. Anna Petrunkina Harrison and Arrayscan experiments were enabled by Veronika Romashova at the JCBC FACS core facilities. Thanks to Liane Dupont and Zheng-Shan Chong for useful advice on the design of CRISPRa screens.

## Author contributions

J.S. performed the arrayed screen. T.W.M.C and A.S.T. performed the CRISPR activation screen. T.W.M.C and A.S.T performed cell binding experiments on RPE1 and CaLu-3 cells. J.S. performed all other binding experiments. I.G. J.A.N. T.P. and A.S.T. expressed and purified S1-Fc fusion protein. J.A.N. supervised the protein purification studies. S.J.W. performed western blotting. B.M.O. performed qPCR. C.M.G-B performed immunostaining and microscopy. J.S. analyzed expression datasets. J.S. analyzed phylogenetic conservation. T.W.M.C. and E.J.D.G. performed RPE1 and CaLu-3 infection experiments. J.S. completed statistical analysis. G.J.W. helped design the study and supervised binding experiments. P.J.L. helped design the study and write the manuscript. J.S. wrote the manuscript with comments from all authors.

## Supplementary information

**Supplemental figure 1.**
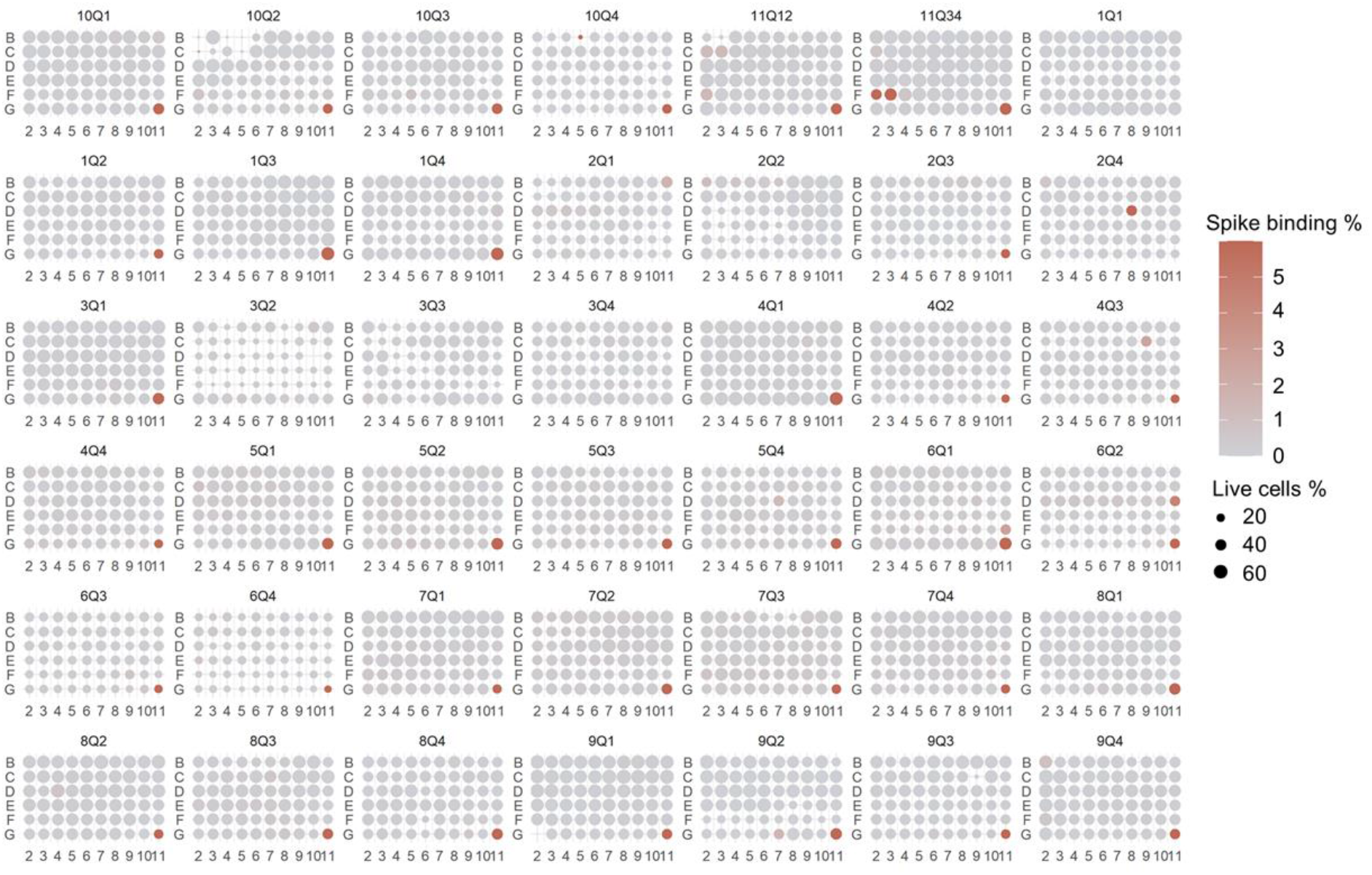
Summary of raw data from arrayed cDNA screen. Each well measured by flow cytometry contains cells transfected with an expression plasmid encoding full-length cDNA from our library encompassing the vast majority of human cell surface receptors. A majority of measured plates included a positive control well of ACE2 in the bottom right corner. As shown in main figure 1C, hits from this screen aside from LRRC15 and already known spike receptors were found to be artifacts upon replication.

**Supplemental figure 2.**
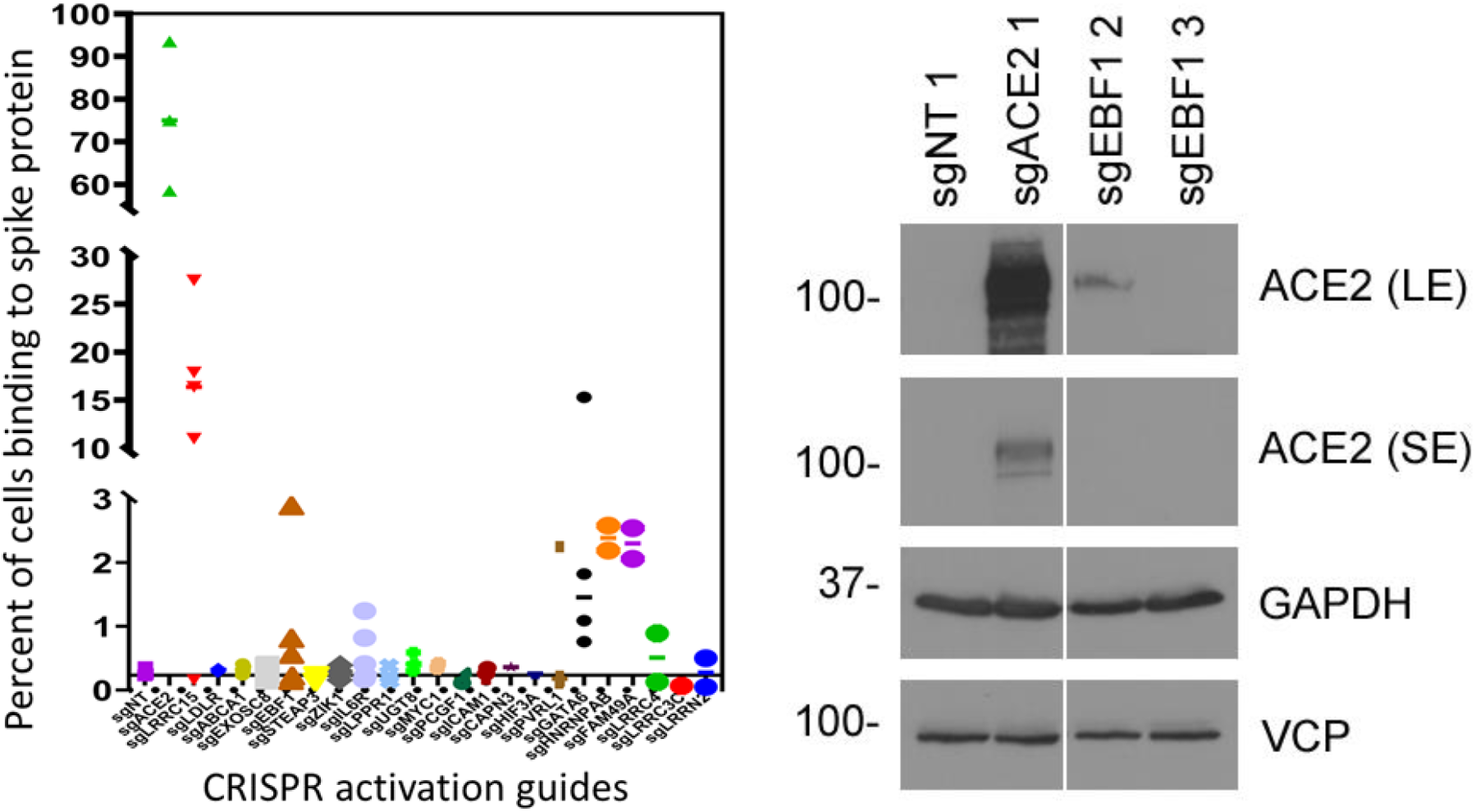
Flow cytometry validation of hits from CRISPRa screen. An overview of binding signals upon replicating each sgRNA is shown (left) alongside western blots that validate one EBF1 sgRNA as slightly upregulating ACE2 expression in RPE1 cells.

**Supplemental figure 3.**
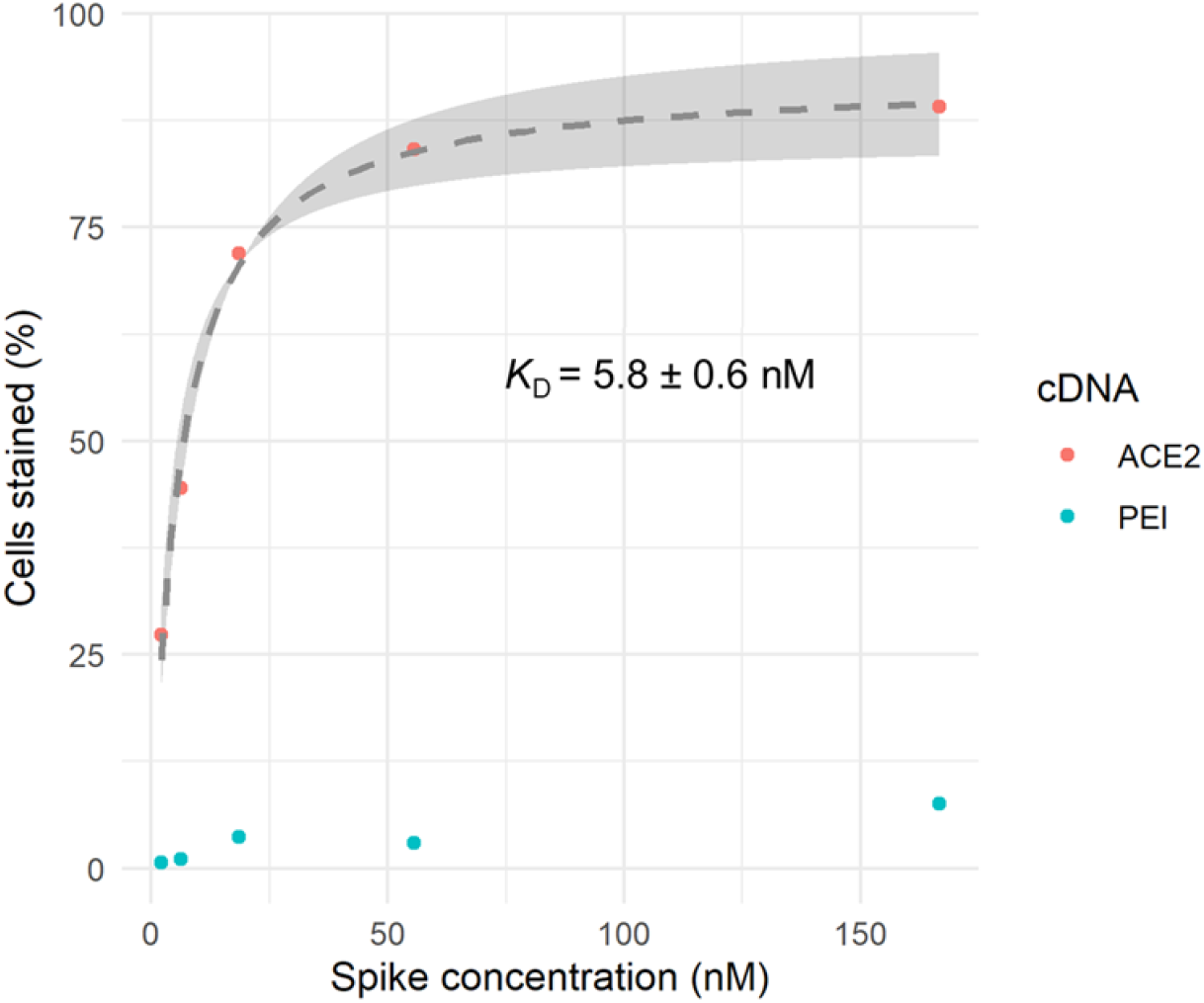
Cell-based spike binding affinity measurements for ACE2 receptor. After incubating ACE2-transfected HEK293 cells with a range of monomeric spike protein concentrations, the binding saturation curve was measured to estimate an equilibrium dissociation constant.

**Supplemental figure 4.**
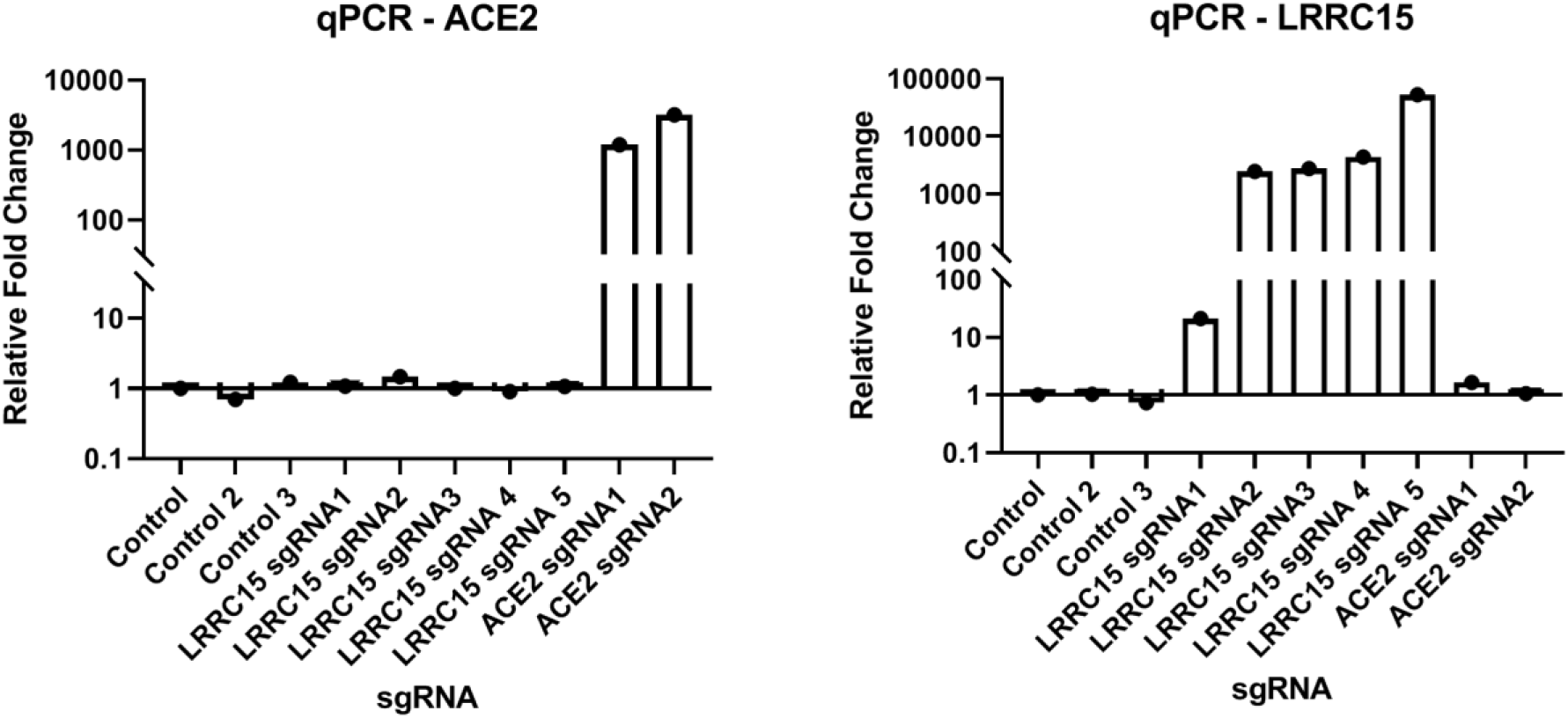
CRISPRa sgRNA targeting *ACE2* or *LRRC15* in RPE-1 SunTag CRISPRa cell line specifically upregulate transcription of the target gene. qPCR measurements of mRNA for *ACE2* or *LRRC15* in cell lines transduced for the sgRNAs indicated along the x-axis.

**Supplemental figure 5.**
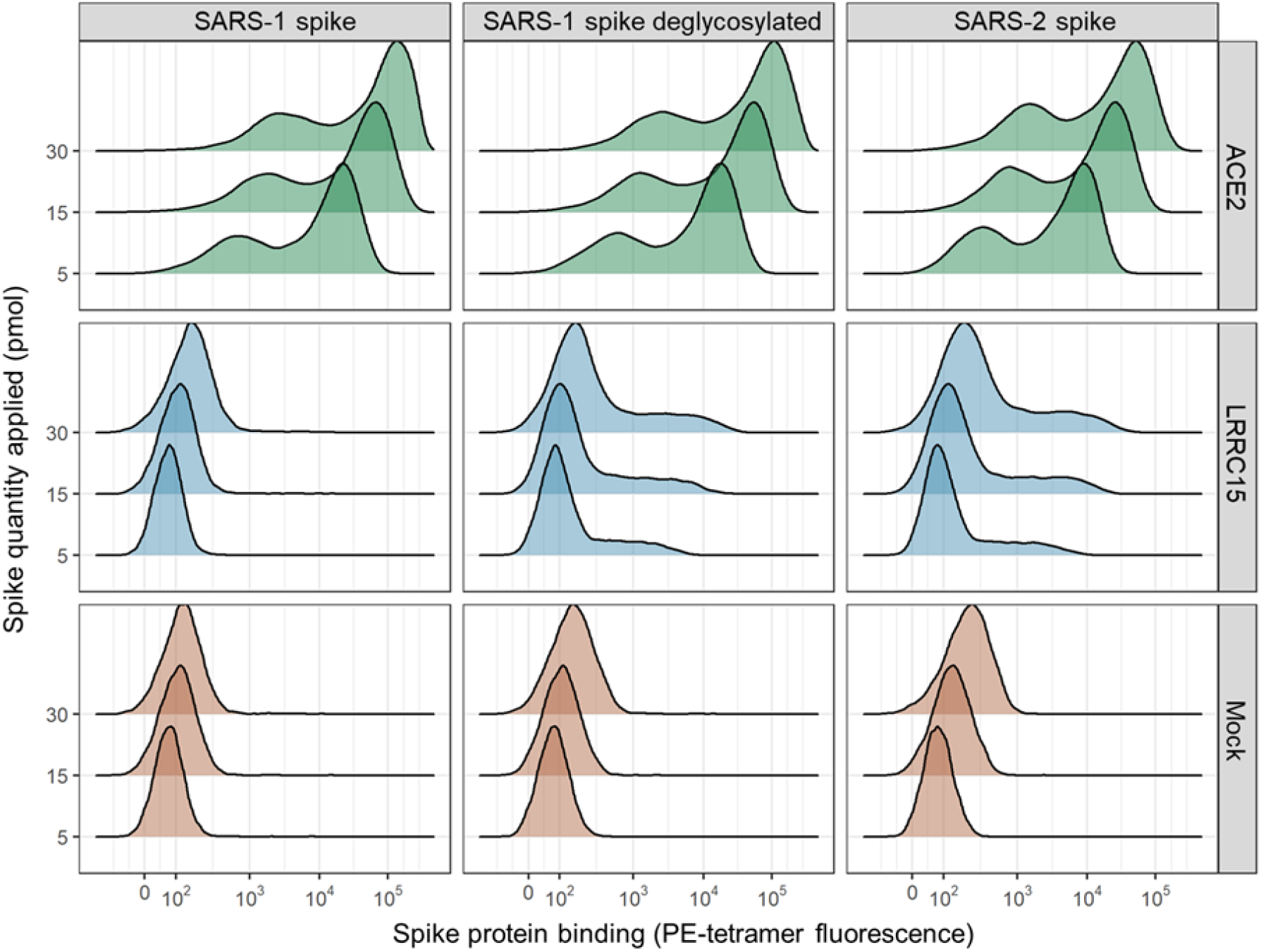
Deglycosylation of SARS-CoV-1 spike restores binding to LRRC15. Comparison of SARS-CoV-1 spike to SARS-CoV-2 spike, where enzymatic removal of most N-linked glycans by PNGase F results in the SARS-CoV-1 spike gaining the ability to bind LRRC15 at similar levels to SARS-CoV-2 spike.

**Supplemental figure 6.**
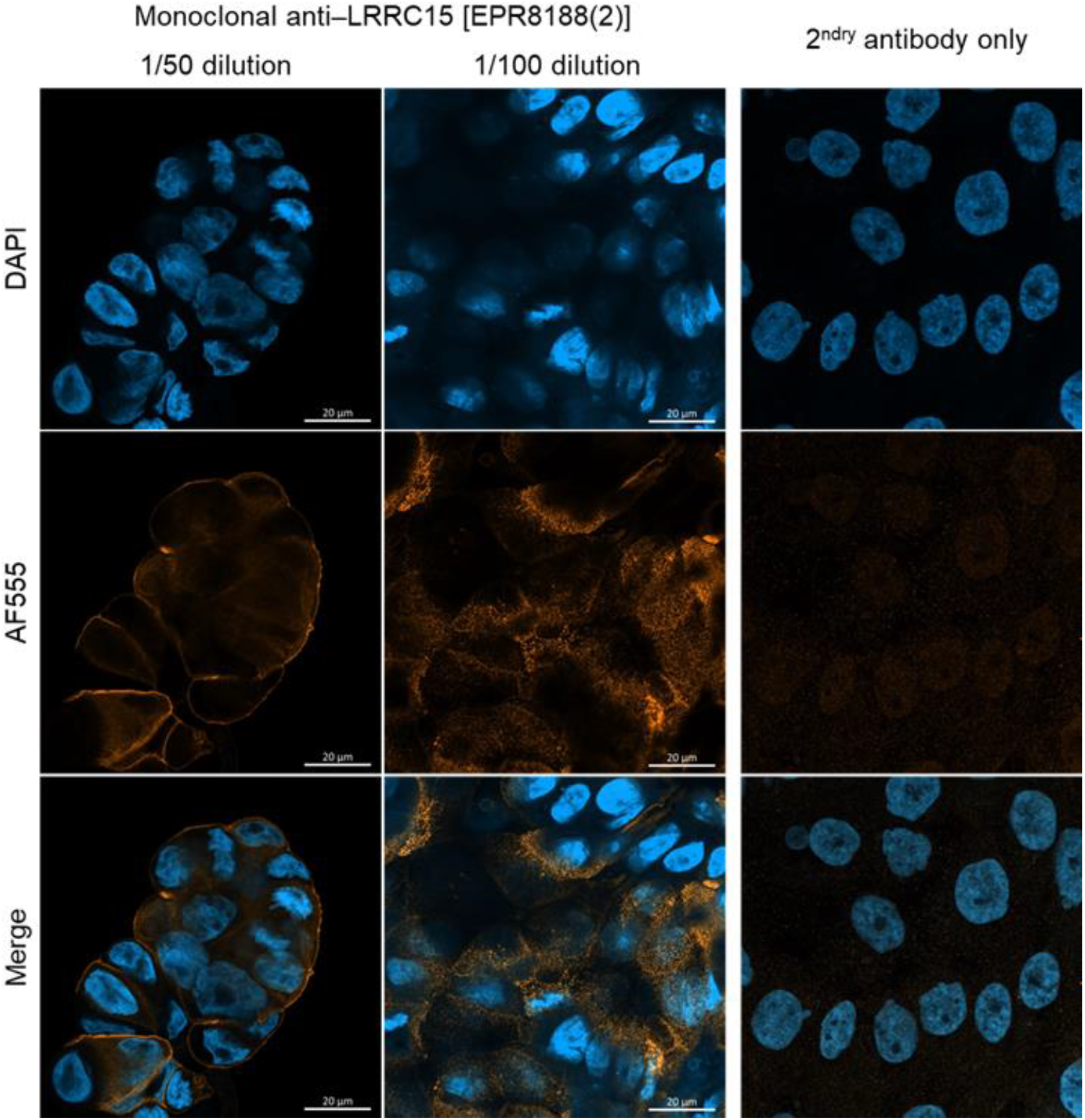
Localization of LRRC15 exogenously expressed in CaLu-3 cells. Two fields of view are shown for CaLu-3 cells stained at different dilutions of monoclonal anti-human LRRC15 antibody. LRRC15 was predominantly detected along the cell plasma membrane, with some possible fainted staining in intracellular compartments.

**Supplemental figure 7.**
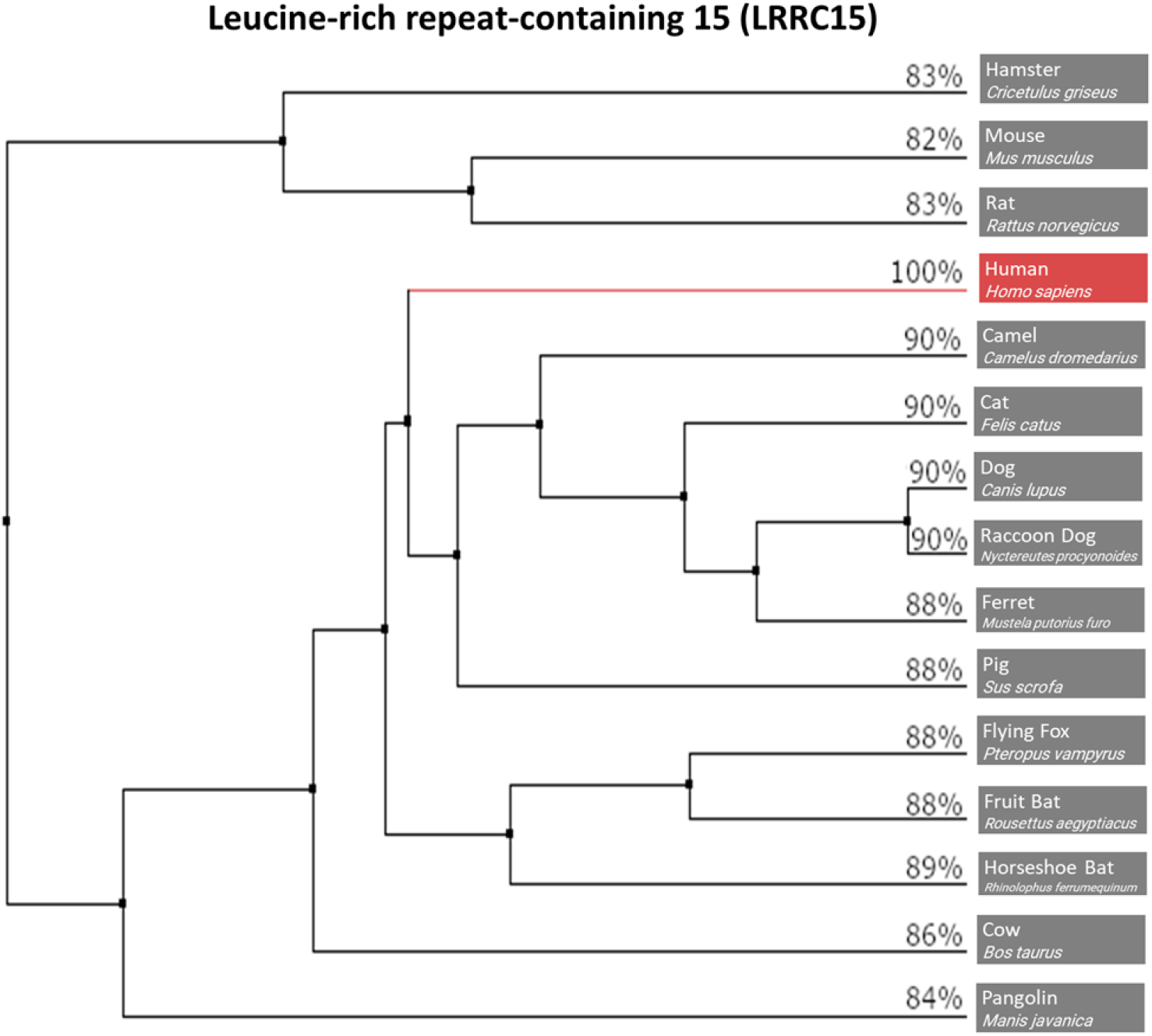
Phylogeny of LRRC15 protein sequence conservation across proposed SARS-CoV-2 hosts and related mammalian species. The percentages of LRRC15 residues that are identical to human are shown at each terminal branch.

**Supplementary data.**
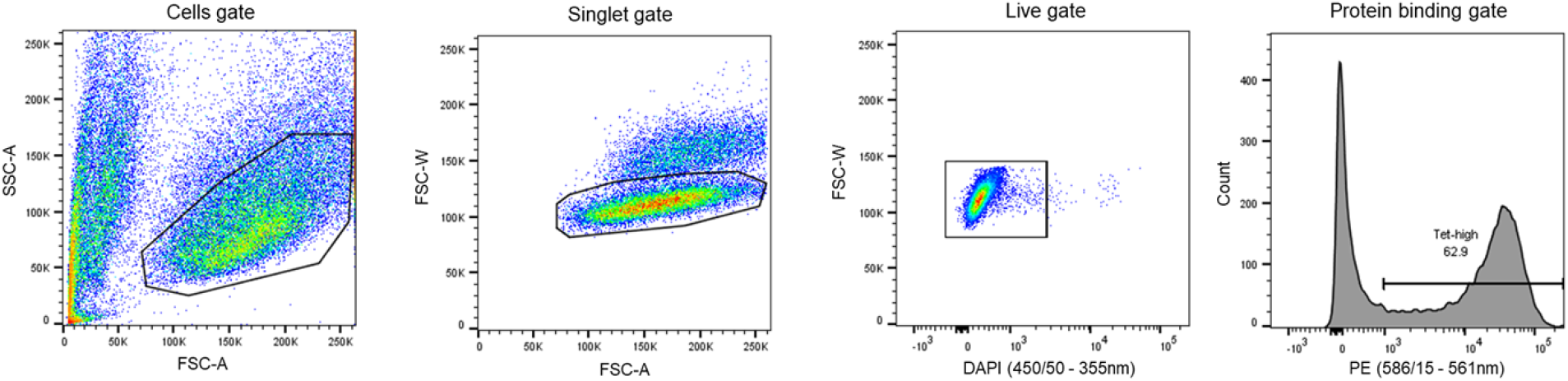
Example gating strategy for cell line flow cytometry measurements of recombinant protein binding. The hierarchy of gates goes from left to right.

